# siDiff: De Novo siRNA Design via Efficacy-Guided Discrete Masked Diffusion

**DOI:** 10.64898/2026.08.11.744092

**Authors:** Zhirun Yue, Honggen Zhang, Xiangrui Gao, Shaokun Shu, Lipeng Lai

**Affiliations:** Peking University Health Science Center; Peking University International Cancer Institute; XtalPi, Inc; Tsinghua University

## Abstract

Target-conditioned *de novo* siRNA design requires capturing both target mRNA binding context and internal siRNA sequence-activity rules. Traditional computational pipelines predominantly rely on discriminative prediction models that rank pre-filtered candidate pools. However, these approaches struggle with generalization on novel target genes and often overlook potent candidates due to dataset size constraints and motif overfitting. To reconcile generation precision with sequence diversity, we propose **siDiff**, an efficacy-guided discrete diffusion framework for target-conditioned *de novo* siRNA design. siDiff pairs a discrete-masked diffusion transformer—which models the underlying sequence distribution over functional duplexes—with a mask-robust efficacy guidance model. During inference, we introduce a biology-aware, three-stage sampling mechanism that performs structural candidate filtering, dynamic unmasking guidance, and cluster-aware redundancy mitigation. This dual mechanism enables the diffusion process to explore broad sequence spaces while the efficacy model prevents distributional shift toward non-functional candidates. Extensive experiments across four datasets, including the public Takayuki benchmark and three curated patent datasets, demonstrate that siDiff significantly outperforms state-of-the-art discriminative baselines and discrete diffusion models, achieving relative hit-rate improvements of over 30% and demonstrating superior generalization on out-of-distribution gene targets. The source code and related materials are available at https://github.com/cybericha/siDiff.

## Introduction

RNA interference (RNAi) technology, powered by small interfering RNAs (siRNAs), has emerged as a cornerstone therapeutic modality for addressing rare genetic disorders and previously undruggable targets. To date, seven siRNA-based therapeutics have received approval from the U.S. Food and Drug Administration (FDA) (Padda et al. 2024). By inducing sequence-specific degradation of complementary messenger RNA (mRNA), siRNAs halt translation and reduce disease-causing protein expression. However, identifying highly potent siRNA candidates remains a costly, labor-intensive, and time-consuming bottleneck, often requiring months of iterative empirical screening across vast candidate pools both *in vitro* and *in vivo*.

Early computational efforts focused on identifying hand-crafted sequence features and thermodynamic parameters from effective siRNAs to score candidate sequences (Khvorova, Reynolds, and Jayasena 2003). As larger standardized databases were constructed (Huesken et al. 2005; Katoh and Suzuki 2007), data-driven approaches emerged, expanding to machine learning regressors, graph neural networks (Bai et al. 2024; Long et al. 2024), and large nucleic acid foundation models (Xiong et al. 2025; Zhang et al. 2025). Despite their success on benchmark datasets, virtually all existing algorithms operate under a rank-based discriminative paradigm (Figure 1, left). These methods evaluate and score a fixed, exhaustively tiled library of candidate sequences against a target mRNA. Consequently, when applied to novel target genes with limited training data, rank-based models frequently struggle to generalize. They tend to overfit to specific training motifs, resulting in poor out-of-distribution performance, overlooked target regions, and missed potent hits.

**Figure 1.**
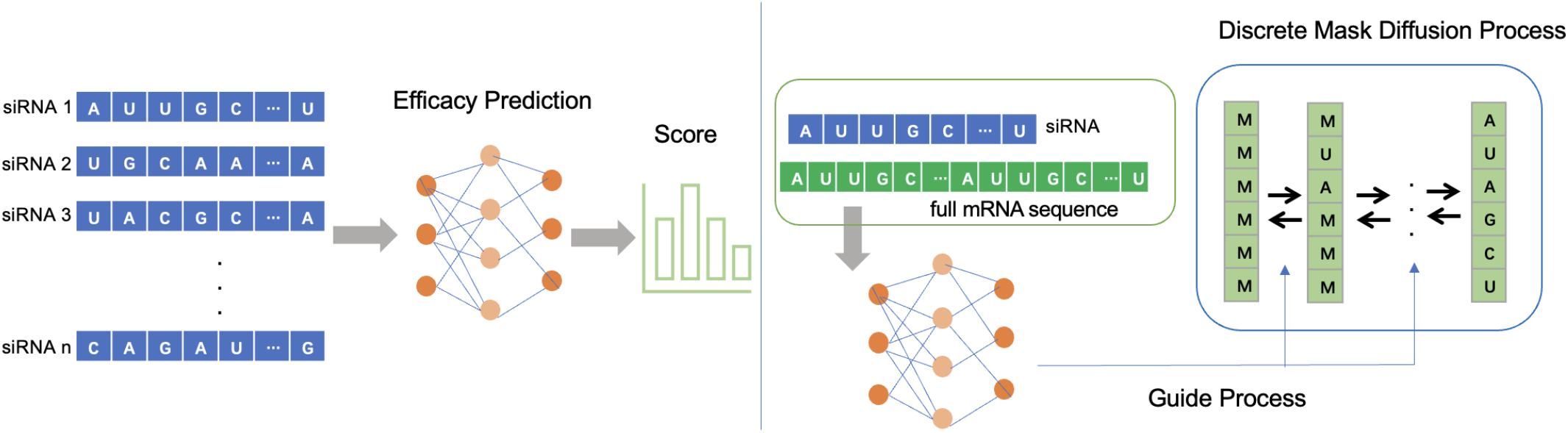
Comparison of siRNA discovery paradigms. **Left:** Traditional rank-based screening pipeline, where a fixed, brute-force candidate library derived from target mRNA is filtered through a prediction model. **Right:** The proposed *de novo* design framework (**siDiff**), which integrates a masked discrete diffusion process with a guided efficacy model to directly generate potent siRNA candidates.

**Figure 2.**
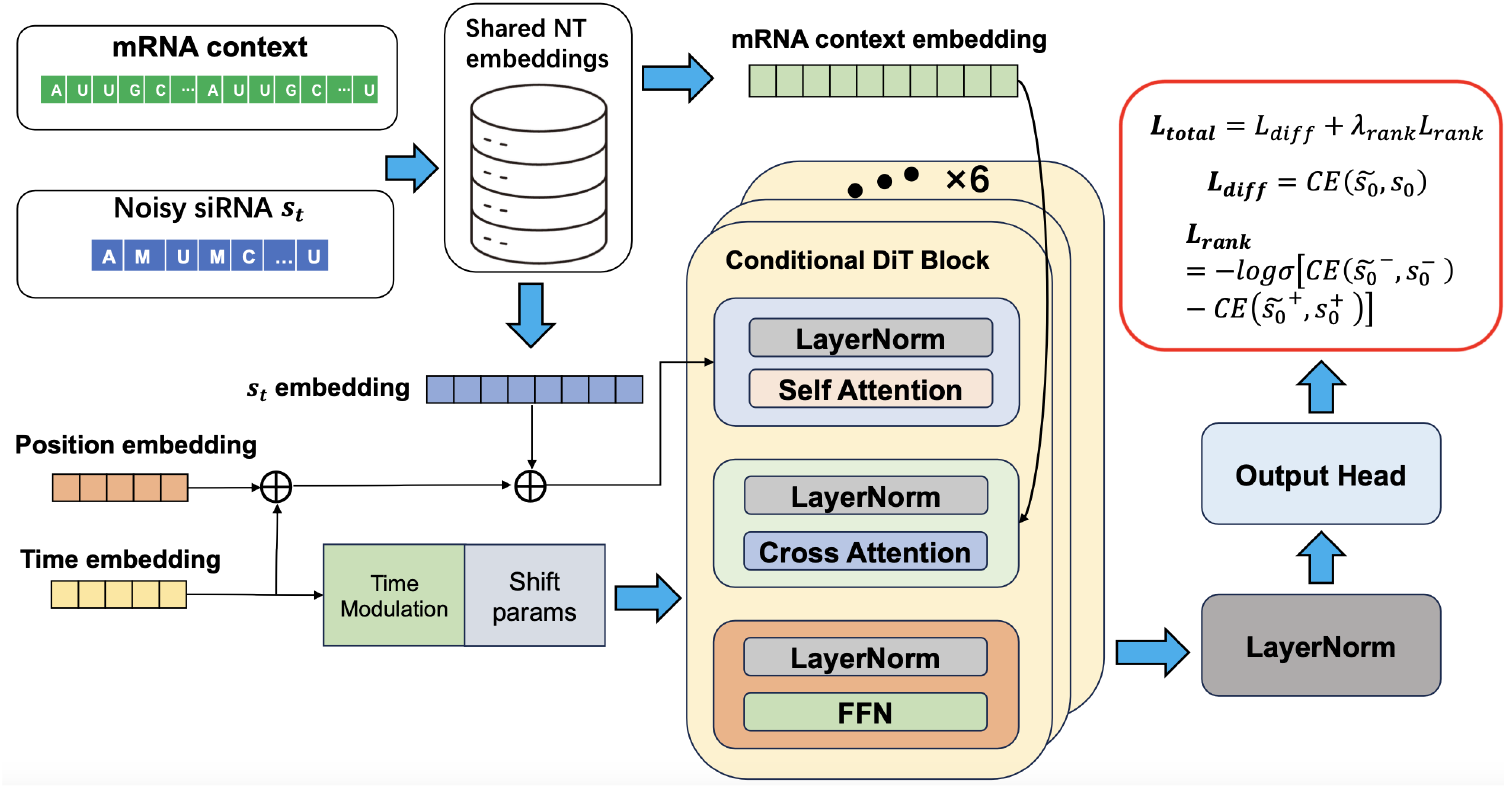
The diagram of the model architecture for training.

To overcome these fundamental limitations, we propose siDiff, a novel *de novo* siRNA design framework that combines masked discrete diffusion with continuous efficacy guidance (Figure 1, right). Rather than exhaustively scoring pre-defined candidate libraries, siDiff models the underlying conditioned distribution of potent siRNA sequences given a target mRNA. During training, we adapt a Discrete Diffusion Transformer architecture (Peebles and Xie 2023) trained on corrupted sequences using an absorbing [`MASK`] state, learning to progressively recover unmasked siRNA duplexes. During inference, we introduce a biology-aware, coarse-to-fine guided sampling strategy executed across three distinct phases:

1. **Structural Constraint & Early Rejection:** The target mRNA is partitioned into overlapping segments to handle sequence length constraints. Starting from a fully masked siRNA candidate (*T* = 21), the generator initiates early unmasking steps (e.g., from *T* = 21 down to *T* = 14). Early non-viable candidates—those failing to form valid reverse-complement duplexes within the target mRNA segment—are filtered out immediately.
2. **Fine-Grained Efficacy-Guided Unmasking:** As sampling progresses to lower noise timesteps, more sequence context is revealed. A pre-trained efficacy predictor dynamically guides the selection of both the next position to unmask (1 ≤ *pos* ≤ 21) and the specific nucleotide assignment (A, U, G, C).
3. **Cluster-Aware Redundancy Mitigation:** Reconstructed active patterns are aligned back to the target mRNA via reverse-complement match to retrieve full siRNA candidates. To prevent spatial clustering of candidates along narrow regions of the mRNA, we apply a cluster-aware diversity mechanism that optimizes both predicted silencing activity and positional sequence diversity.

We evaluate siDiff across four benchmark datasets, including the public Takayuki (Katoh) dataset (Katoh and Suzuki 2007) and three patent-derived datasets. To rigorously benchmark *de novo* sequence generation quality, we formulate generation accuracy metrics (Hit_*δ*_@*K*) alongside structural diversity metrics. Experimental results demonstrate that siDiff significantly outperforms state-of-the-art rank-based discriminative baselines across all datasets.

Our primary contributions are summarized as follows:

- We introduce **siDiff**, shifting siRNA discovery from traditional discriminative scoring to a target-conditioned *de novo* generative diffusion paradigm.
- We propose a biology-aware, coarse-to-fine guided sampling method that couples a discrete masked diffusion transformer with a fine-grained efficacy model to steer sequence generation toward potent candidates.
- We define a new evaluation framework and metrics specifically tailored for generative siRNA models, establishing benchmarks for generation accuracy and sequence diversity.
- Extensive empirical evaluations show that siDiff achieves state-of-the-art performance, outperforming strong baselines with relative hit-rate improvements of approximately 35% on the Takayuki dataset and 38% on the CTNNB1 target dataset, respectively.

## Related Work

### Discrete Diffusion Models for Sequence Generation

Diffusion models have achieved remarkable success in continuous domains such as image and video synthesis (Ho, Jain, and Abbeel 2020; Rombach et al. 2022; Ho et al. 2022; Peebles and Xie 2023). However, their direct application to discrete modalities, such as text and biological sequences, has historically been hindered by the quantization errors inherent in mapping categorical tokens to continuous latent spaces (Li et al. 2022). To address this limitation, Discrete Denoising Diffusion Probabilistic Models (DDDPMs) introduced discrete Markov transition matrices over categorical distributions (Austin et al. 2021). Building upon this foundation, Masked Diffusion Language Models (MDLMs) simplified absorbing-state diffusion losses into classical masked modeling objectives via substitution-based parameterizations (Sahoo et al. 2024). Recent large-scale discrete diffusion frameworks have further demonstrated the advantages of non-autoregressive, bidirectional generation over traditional autoregressive paradigms (Li et al. 2026). In these architectures, the forward process progressively corrupts tokens into an absorbing [MASK] state, while the reverse process learns to globally unmask sequences, enabling parallel and flexible conditional generation.

### siRNA Design and Efficacy Prediction

Early approaches to siRNA efficacy prediction relied heavily on handcrafted biological features (Naito et al. 2009; Khvorova, Reynolds, and Jayasena 2003) and empirical rules (Katoh and Suzuki 2007). The curation of large-scale standardized naked siRNA datasets (Huesken et al. 2005) enabled data driven based approaches, ranging from linear regression frameworks such as DSIR and i-Score to support vector machines (Wang, Huang, and Yang 2010) and ensemble methods (Monopoli, Korkin, and Khvorova 2023). Subsequent advances in deep learning significantly improved predictive capacity through convolutional architectures (Han et al. 2018), graph neural networks (La Rosa et al. 2022; Long et al. 2024), and latent representation models (He et al. 2017; Bereczki et al. 2025). Most recently, Transformer-based architectures have emerged as the state of the art for modeling complex sequence-activity relationships in small RNA molecules (Liu et al. 2024; Bai et al. 2024; Zhang, Gao, and Lai 2025). Despite these predictive advances, the problem of guided de novo generation of optimized siRNA sequences remains a challenging frontier, motivating the integration of discrete diffusion models into siRNA drug discovery workflows.

### Problem Statement

We study target-conditioned *de novo* siRNA design. Let **m** = (*m*_1_, …, *m*_*L*_) ∈ *A*^*L*^ denote a target mRNA sequence of length *L*, and let 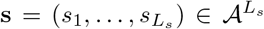 denote a 21-nucleotide siRNA sequence (*L*_*s*_ = 21), where *A* = {A, C, G, U}. We model the conditional distribution *p*_*θ*_(**s | m**) to capture the complex sequence-dependent interactions between target mRNA regions and functional siRNA candidates. Given an input mRNA **m**, our goal is to learn a conditional generative process that directly generates diverse, site-specific siRNAs exhibiting high knockdown efficiency.

### Methodology

We propose **siDiff**, an efficacy-guided generative framework for *de novo* siRNA design. As illustrated in Figure 1, siDiff integrates three core components: (i) a conditional discrete masked diffusion model trained on highly effective mRNA– siRNA pairs, (ii) a mask-robust efficacy guidance model, and (iii) a biology-aware structured sampling pipeline.

Specifically, the masked diffusion model establishes a conditional generative prior over potent sequence motifs. In parallel, the mask-robust guidance model predicts the latent knockdown efficiency of partially masked sequence states during reverse diffusion, supplying dynamic gradients that steer sampling trajectories toward potent candidates. During inference, our biology-aware sampling pipeline enforces spatial search-space diversity while applying thermodynamic and reverse-complement constraints to produce site-specific, highly potent siRNAs.

### Masked Discrete Diffusion siRNA Model

We formulate target-conditioned siRNA generation as a discrete masked diffusion process. Given a target mRNA **m** and its corresponding siRNA sequence 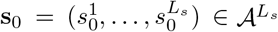, the forward corruption process progressively substitutes nucleotides with an absorbing [MASK] token. Let *A*_M_ = *A* ∪ [MASK] represent the augmented vocabulary.

Following an absorbing-state Markov transition framework (Austin et al. 2021; Sahoo et al. 2024), the probability of a nucleotide remaining unmasked at timestep *t* ∈ {1, …, *T*} is defined as 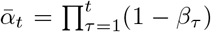, where *β*_*τ*_ ∈ (0, 1) governs the noise schedule. The forward marginal distribution for position *i* ∈ {1, …, *L*_*s*_} is expressed as:

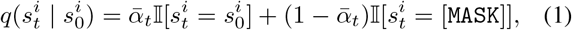

where I[·] is the indicator function. Under this formulation, once a nucleotide transitions into the absorbing [MASK] state, it remains masked until *t* = *T*, where 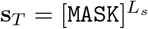.

The reverse denoising process aims to recover the original siRNA sequence **s**_0_ conditioned on the target mRNA context **m**:

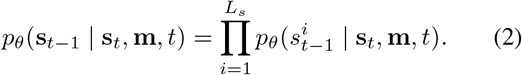

Generation initializes from a fully masked state **s**_*T*_ and iteratively predicts unmasked token assignments across timesteps guided by the conditional distribution learned by the network.

#### Training and Context Augmentation

To bias the generative prior toward highly potent candidates, we train the discrete diffusion model exclusively on experimentally validated mRNA–siRNA pairs whose inhibition levels exceed a predefined activity threshold *τ*_eff_ :

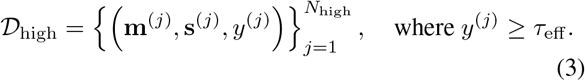

Here, **m**^(*j*)^ represents the mRNA context, **s**^(*j*)^ is the validated 21-nt siRNA duplex, and *y*^(*j*)^ is the experimentally measured inhibition efficiency.

#### Random-Window Context Augmentation

Naïvely training on fixed mRNA context windows can lead the diffusion transformer to exploit positional shortcuts (e.g., memorizing target sites at fixed offsets). To force the model to learn true sequence-dependent mRNA–siRNA binding dynamics, we implement a random-window context augmentation strategy during training.

For a total context window size of *L*_*m*_, we extract sub-segments surrounding the *L*_*s*_-nt target site within the full-length target transcript **M**. We randomly sample an upstream flank length *ℓ >* 0 and compute the corresponding down-stream flank length *r >* 0 as:

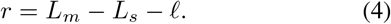

The augmented mRNA context vector **m**^(*ℓ*)^ is then defined as:

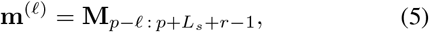

where *p* denotes the start index of the target binding site in **M**. By dynamically altering *ℓ* across training epochs, the target site appears at varying relative positions within **m**^(*ℓ*)^, encouraging invariant learning of target-site complementary relationships.

#### Training Objective

Each training instance is represented as a tuple (**m, s**_0_, *y*) ∈ *D*_high_, where **s**_0_ denotes the uncorrupted 21-nt siRNA sequence, **m** is the corresponding target mRNA context, and *y* is the experimentally measured inhibition efficiency. During training, a diffusion timestep *t* is sampled uniformly from *U* (1, *T*), and a corrupted state **s**_*t*_ ~ *q*(**s**_*t*_ | **s**_0_) is generated via the forward masking process (Eq. 1). Let 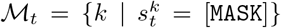 denote the set of indices corresponding to masked nucleotide positions.

#### Masked Sequence Reconstruction Loss

The primary generative training objective is the masked-token cross-entropy loss, evaluated strictly over the masked positions *M*_*t*_:

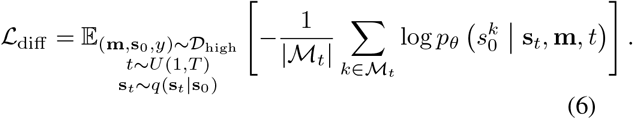

This objective forces the Conditional Diffusion Transformer (CondDiT_*θ*_) to learn target-conditioned contextual rules for recovering unmasked nucleotides from partially observed sequence states.

#### Auxiliary Efficacy-Aware Ranking Loss

Although samples in *D*_high_ satisfy the efficacy threshold *τ*_eff_, their measured inhibition values provide additional supervision about relative potency. We therefore apply an auxiliary pairwise ranking objective to clean sequences (*t* = 0), avoiding noise introduced by diffusion corruption. For each sample (**m**^(*i*)^, 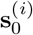, *y*^(*i*)^), the clean CondDiT representation is pooled and projected by a linear ranking head to obtain a scalar score *r*^(*i*)^.

To reduce sensitivity to experimental fluctuations, we retain only pairs whose efficacy difference exceeds a margin *δ*:

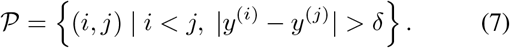

Let *z*_*ij*_ = sign(*y*^(*i*)^ − *y*^(*j*)^). The ranking loss is

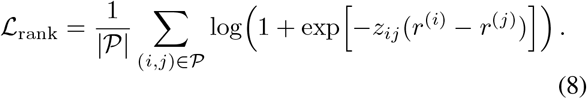

#### Joint Training Objective

The unified objective optimizes token recovery while simultaneously organizing the latent representation space by inhibitory efficiency:

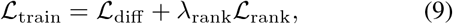

where *λ*_rank_ ≥ 0 is a hyperparameter balancing reconstruction fidelity and rank-ordering alignment.

### Biology-Aware Sampling Strategy

Navigating the generation process within the sparse high-efficacy siRNA manifold poses a critical challenge for pre-trained diffusion models in ensuring both candidate diversity and predictive accuracy. To address this, our biology-aware sampling strategy couples the generative prior with a pre-trained efficacy predictor, leveraging bio-domain rules to effectively constrain the sampling dynamics within the target functional manifold.

#### Algorithm 1

Coarse-to-Fine Efficacy-Guided Sampling

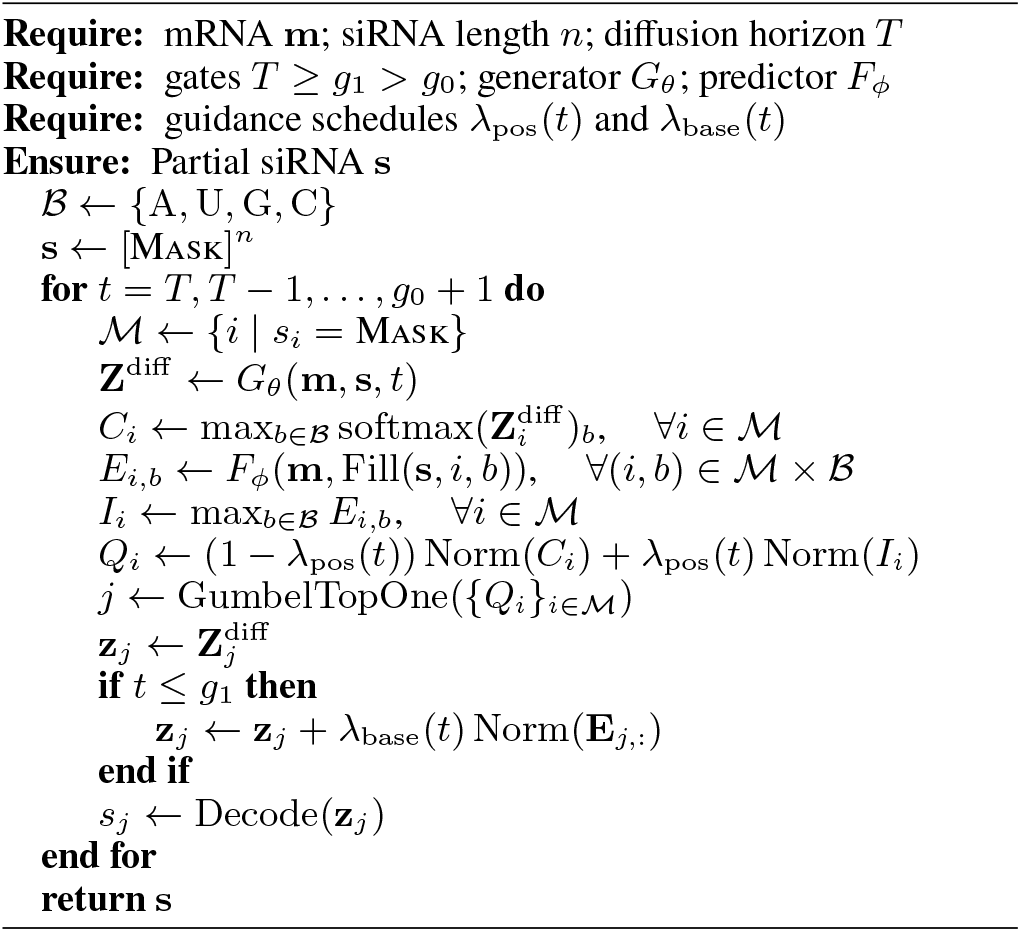

The position-level efficacy guidance is active throughout the sampling trajectory and is controlled by *λ*_pos_(*t*). In contrast, nucleotide-level guidance is delayed:

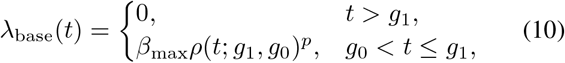

where *ρ*(*t*; *g*_1_, *g*_0_) is a monotonic progress function over the base-guidance phase, *β*_max_ controls the maximum guidance strength, and *p* determines the shape of the schedule.

#### Efficacy-Guided Active Pattern Proposal

Generating a complete siRNA from a fully masked state involves substantial uncertainty. We therefore combine the sequence prior of the diffusion model with efficacy estimates from a predictor trained on partially masked sequences. This coarse-to-fine strategy progressively guides sampling toward high-efficacy regions.

An active pattern is a partial siRNA containing efficacy-relevant nucleotide assignments identified during guided diffusion. We use *g*_0_ to denote the terminal boundary of efficacy-guided active-pattern sampling: denoising is performed only for *t > g*_0_, and the partial sequence obtained at this boundary is retained as the active-pattern proposal.

To improve the discovery of informative active patterns, the position selected at each denoising step should reflect the generator’s confidence along the positional dimension. For each unresolved position *i* ∈ *U*_*t*_, we define the generation confidence 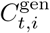 as the maximum probability assigned by the generator to the four possible nucleotides. However, relying solely on this confidence may repeatedly favor positions that are already dominant under the learned generative distribution, causing premature concentration of the sampling trajectories and limiting the diversity of the resulting active-pattern proposals.

We therefore use the efficacy predictor to dynamically correct the position confidence. Specifically, the predictor evaluates the four hypothetical nucleotide assignments at each unresolved position, and their maximum predicted efficacy is used as the position-level efficacy potential 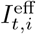. Because generation confidence and efficacy potential have different numerical scales, they are independently normalized over the current unresolved-position set *U*_*t*_. The modified confidence is defined as

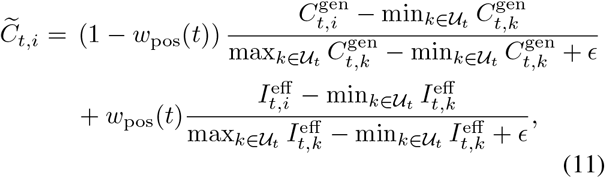

where *w*_pos_(*t*) is a timestep-dependent guidance weight. This formulation preserves the generator’s learned sequence prior while progressively increasing the preference for positions with greater predicted efficacy potential.

To retain stochastic exploration, Gumbel noise is added to the temperature-scaled modified confidence before selecting the Top-1 position. The Gumbel distribution is particularly suitable for this operation because it characterizes the limiting distribution of extrema and enables stochastic categorical selection through the Gumbel-Max principle. The overall position sampler is abstracted as

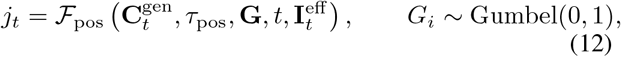

where *τ*_pos_ controls the exploration strength and **G** provides the random perturbations used for stochastic Top-1 selection.

We divide nucleotide-type sampling into two phases at *g*_1_, since the reliability of nucleotide-level efficacy estimates depends on the completeness of the partially generated sequence. In the early phase (*t > g*_1_), a large fraction of nucleotides remains masked, making the efficacy difference among candidate bases noisy and potentially misleading.

Therefore, after selecting the position to be unmasked, its nucleotide type is sampled solely from the diffusion-model logits. In the later phase (*g*_0_ *< t* ≤ *g*_1_), the partial sequence provides a more informative context, and nucleotide-level efficacy is incorporated to favor bases that are predicted to improve silencing efficacy.

Specifically, for the selected position *j*, we substitute each candidate *b* ∈ {A, U, G, C} and evaluate

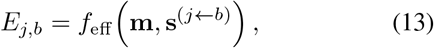

where **m** denotes the target mRNA context and **s**^(*j*←*b*)^ is the current partial sequence with position *j* temporarily filled by *b*. The four efficacy scores are standardized as

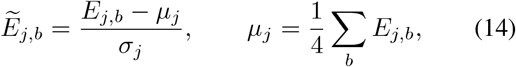

where *σ*_*j*_ is the standard deviation over the four candidate bases. If *σ*_*j*_ is numerically negligible, we set 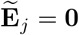.

The efficacy-guided nucleotide logits are then computed by

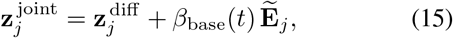

where 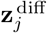 denotes the diffusion logits over the four nucleotide types. We gradually increase the guidance strength according to

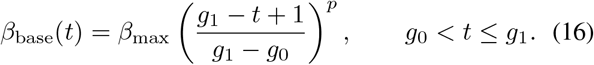

Finally, the nucleotide is sampled from the temperature-scaled distribution 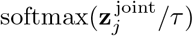 (or selected by its maximum in greedy decoding). The details can be found in Algorithm 1.

#### Candidate Derivation and Clustering

Because an active pattern specifies only a subset of siRNA positions, it can match multiple regions of the target mRNA. We align its instantiated nucleotides to the mRNA under reverse-complementarity and a predefined mismatch tolerance, while treating masked positions as unconstrained. Each match is expanded into a valid 21-nt target window, yielding a one-to-many mapping from an active pattern to complete siRNA candidates.

To reduce redundancy among nearby or overlapping windows, candidates are sorted by their start positions and greedily clustered. A candidate with start *x* is added to the current cluster *C* if

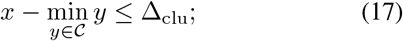

otherwise, it starts a new cluster. Bounding the total cluster span preserves local alternatives while preventing spatially distant candidates from being merged through transitive chaining.

#### Inhibition-Aware Candidate Evaluation and Sampling

The active pattern identifies promising target regions but does not represent the final siRNA. For each derived 21-nt mRNA target site, we construct its reverse-complementary siRNA and retrieve the corresponding upstream and downstream mRNA contexts. An inhibition predictor then assigns an efficacy score *q*_*a*_ to each completed candidate. This provides deterministic efficacy assessment while retaining candidates from multiple target regions for diverse generation.

Because neighboring target sites may exhibit correlated efficacy (Davis et al. 2025), each cluster *G*_*k*_ is summarized by its mean score and high-efficacy ratio:

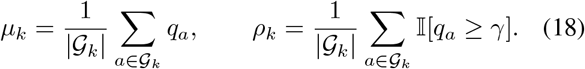

We then apply three-grade staircase sampling. Clusters satisfying progressively relaxed thresholds on *µ*_*k*_ and *ρ*_*k*_ receive sampling quotas *K*_1_ *> K*_2_ *> K*_3_, while clusters failing the final grade are discarded. Within each retained cluster, candidates are ranked by *q*_*a*_ and sampled up to the assigned quota. This cluster-aware strategy balances inhibition efficacy with coverage across distinct mRNA regions.

## Experiments

### Experimental Setup

#### Datasets

We integrate experimentally measured siRNAs from the Huesken dataset, the Takayuki dataset, and three patent datasets targeting ANGPTL7, CTNNB1, and GSK3A. We standardize all sequences and min-max normalize inhibition values within each mRNA. We retain high-activity sites with normalized inhibition ≥ 0.7. For each retained site, we sample up to five 61-nt windows from the full mRNA using randomly selected valid upstream and downstream offsets, while ensuring that the complete 21-nt target site is retained in each window. Table 1 summarizes the numbers of raw measurements, retained relatively high-activity sites, and constructed context windows for each dataset. The models are trained on Huesken-derived data, whereas Takayuki and the patent-derived datasets are excluded from training and used for external evaluation.

**Table 1:** Statistics of the de novo siRNA design dataset.

| Dataset | Raw records | Active sites | Context views |
| --- | --- | --- | --- |
| Takayuki | 702 | 190 | 943 |
| ANGPTL7 | 111 | 31 | 155 |
| CTNNB1 | 330 | 242 | 1210 |
| GSK3A | 122 | 78 | 390 |
| Total | 1265 | 541 | 2698 |

#### Baselines

We compare siDiff against five baselines. *Random Sampling* uniformly samples valid target starts from the 61-nt context. *Enumeration + iScore* and *Enumeration + OligoFormer* enumerate all 41 valid starts, construct the corresponding reverse-complement siRNAs, and rank them using iScore and OligoFormer, respectively. *Vanilla Diffusion* uses the same CrossAttentionDiT generator as siDiff but removes efficacy guidance and predictor-based selection. *Guided Diffusion + Efficacy Selection* applies efficacy guidance during diffusion and directly selects generated candidates according to their predicted efficacy.

#### Evaluation protocol

Given a 61-nt mRNA context without its target position, each method returns a ranked list of candidate starts. Position recovery is measured by

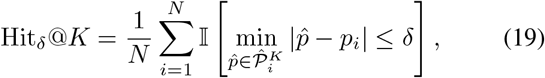

where 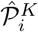 contains the top-*K* unique predicted starts and *p*_*i*_ is the annotated high-activity start. We report Hit_0_@1 and Hit_3_@3.

We further introduce Efficacy-Window Coverage (EWC) to measure the recovery of diverse high-efficacy regions. Each mRNA is partitioned into non-overlapping 100-nt windows. For each window *B*_*j*_, we compute the number of measured sites *o*_*j*_, the proportion of high-efficacy sites *r*_*j*_, and their average efficacy *a*_*j*_. The ground-truth efficacy-window set is

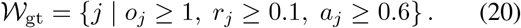

After deduplicating generated target sites, we construct *W*_pred_ analogously using their predicted efficacy scores. EWC is defined as

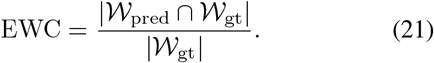

All metrics are computed and reported separately for each mRNA. The resulting | *W*_gt_ | values are 1, 1, 29, and 9 for Takayuki, ANGPTL7, CTNNB1, and GSK3A, respectively. For Takayuki, the high-efficacy sites are strongly concentrated within one region. In contrast, ANGPTL7 contains only 31 experimentally measured high-efficacy sites, leaving only one window that satisfies the efficacy-window criteria. Detailed metric definitions and implementation protocols are provided in the supplementary material.

#### Implementation details

The generator is built upon a Diffusion Transformer (DiT), with mRNA conditions incorporated through cross-attention. The efficacy guidance model adopts a Transformer-based encoder–decoder architecture to estimate inhibition for partially masked siRNA–mRNA pairs. We set the weight of the auxiliary ranking loss to *λ*_rank_ = 0.2. All experiments are conducted on an NVIDIA A100 GPU.

### Partial-State Prediction and Target-Site Recovery

We evaluate curriculum-based efficacy prediction on partially masked siRNAs using Pearson (PCC) and Spearman (SCC) correlation coefficients. As shown in Table 2, the PCC and SCC results confirm reliable efficacy prediction under partial masking.

**Table 2:** Validation performance of the efficacy predictor across curriculum stages.

| Curriculum stage | Val. PCC ↑ | Val. SCC ↑ |
| --- | --- | --- |
| Stage 1 | 0.643 | 0.641 |
| Stage 2 | 0.662 | 0.668 |
| Stage 3 | 0.677 | 0.684 |

Given the clustered candidate-start set *C*_*i*_ and annotated high-activity start *p*_*i*_ for context *i*, we measure candidate-pool recovery at positional tolerance *δ* as

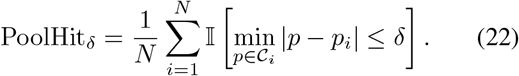

Table 3 reports PoolHit_*δ*_ for *δ* ∈ {0, 1, 2}, showing that active-pattern candidates preserve annotated target regions and generalize beyond the training distribution.

**Table 3:** Target-site coverage of the clustered candidate pools.

| Dataset | PoolHit<sub>0</sub> ↑ | PoolHit<sub>1</sub> ↑ | PoolHit<sub>2</sub> ↑ |
| --- | --- | --- | --- |
| Takayuki | 0.78 | 0.96 | 0.97 |
| ANGPTL7 | 0.84 | 0.89 | 0.93 |
| CTNNB1 | 0.72 | 0.84 | 0.92 |
| GSK3A | 0.60 | 0.92 | 0.94 |

### Efficacy-Guided Design with Diversity Preservation

Table 4 compares generation performance across four external datasets. siDiff achieves the best Hit_0_@1 and Hit_3_@3 on every dataset while recovering all ground-truth efficacy windows except one on CTNNB1. Compared with enumeration-based methods, siDiff improves Hit_0_@1*/*Hit_3_@3 from 0.038*/*0.401 (iScore) and 0.048*/*0.318 (OligoFormer) to 0.242*/*0.664 on Takayuki, and from 0.022*/*0.385 and 0.023*/*0.275 to 0.200*/*0.654 on CTNNB1. Vanilla Diffusion and Guided Diffusion + Efficacy Selection also attain competitive EWC, reaching 25*/*29 and 26*/*29 on CTNNB1 and 7*/*9 and 8*/*9 on GSK3A, respectively. However, their Hit_3_@3 scores remain substantially below those of siDiff. Overall, siDiff consistently combines accurate target-site recovery with broad efficacy-window coverage, demonstrating strong generalization across all four external datasets.

**Table 4:**
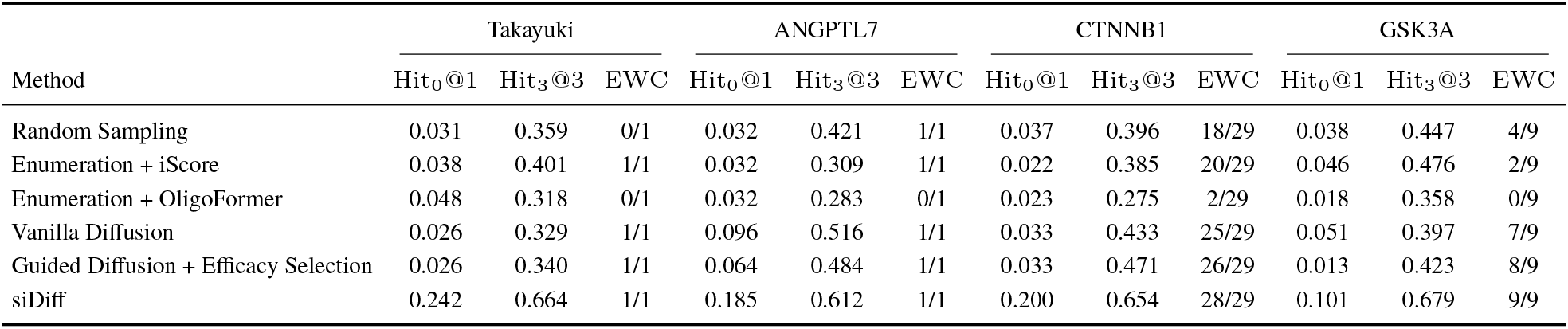
Generation performance on four evaluation datasets.

| Method | Takayuki |  |  | ANGPTL7 |  |  | CTNNB1 |  |  | GSK3A |  |  |
| --- | --- | --- | --- | --- | --- | --- | --- | --- | --- | --- | --- | --- |
|  | Hit<sub>0</sub>@1 | Hit<sub>3</sub>@3 | EWC | Hit<sub>0</sub>@1 | Hit<sub>3</sub>@3 | EWC | Hit<sub>0</sub>@1 | Hit<sub>3</sub>@3 | EWC | Hit<sub>0</sub>@1 | Hit<sub>3</sub>@3 | EWC |
| Random Sampling | 0.031 | 0.359 | 0/1 | 0.032 | 0.421 | 1/1 | 0.037 | 0.396 | 18/29 | 0.038 | 0.447 | 4/9 |
| Enumeration + iScore | 0.038 | 0.401 | 1/1 | 0.032 | 0.309 | 1/1 | 0.022 | 0.385 | 20/29 | 0.046 | 0.476 | 2/9 |
| Enumeration + OligoFormer | 0.048 | 0.318 | 0/1 | 0.032 | 0.283 | 0/1 | 0.023 | 0.275 | 2/29 | 0.018 | 0.358 | 0/9 |
| Vanilla Diffusion | 0.026 | 0.329 | 1/1 | 0.096 | 0.516 | 1/1 | 0.033 | 0.433 | 25/29 | 0.051 | 0.397 | 7/9 |
| Guided Diffusion + Efficacy Selection | 0.026 | 0.340 | 1/1 | 0.064 | 0.484 | 1/1 | 0.033 | 0.471 | 26/29 | 0.013 | 0.423 | 8/9 |
| siDiff | 0.242 | 0.664 | 1/1 | 0.185 | 0.612 | 1/1 | 0.200 | 0.654 | 28/29 | 0.101 | 0.679 | 9/9 |

### Ablation Study

Table 5 reports ablations of the guidance and candidate-selection components on four external benchmarks. Relative to no guidance, position-only and base-only guidance improve the mean Hit_3_@3 by 0.179 and 0.174, respectively, while also increasing EWC on the multi-window datasets. Full-Trajectory Guided Diffusion recovers 25*/*29 efficacy windows on CTNNB1 and 6*/*9 on GSK3A despite lower position accuracy, suggesting that diffusion sampling preserves diversity and supports exploratory target-site discovery. Compared with siDiff w/o clustering, clustering improves the mean Hit_0_@1 and Hit_3_@3 by 0.150 and 0.183, while recovering seven and three additional windows on CTNNB1 and GSK3A, respectively. Clustering is therefore critical for effective target-site proposal.

**Table 5:** Ablation study of sampling guidance components on four evaluation datasets.

| Variant | Takayuki |  |  | ANGPTL7 |  |  | CTNNB1 |  |  | GSK3A |  |  |
| --- | --- | --- | --- | --- | --- | --- | --- | --- | --- | --- | --- | --- |
|  | Hit<sub>0</sub>@1 | Hit<sub>3</sub>@3 | EWC | Hit<sub>0</sub>@1 | Hit<sub>3</sub>@3 | EWC | Hit<sub>0</sub>@1 | Hit<sub>3</sub>@3 | EWC | Hit<sub>0</sub>@1 | Hit<sub>3</sub>@3 | EWC |
| No guidance | 0.104 | 0.487 | 1/1 | 0.193 | 0.417 | 1/1 | 0.221 | 0.442 | 11/29 | 0.166 | 0.487 | 2/9 |
| Position-only guidance | 0.131 | 0.639 | 1/1 | 0.193 | 0.612 | 1/1 | 0.186 | 0.631 | 20/29 | 0.154 | 0.666 | 7/9 |
| Base-only guidance | 0.140 | 0.628 | 1/1 | 0.193 | 0.661 | 1/1 | 0.173 | 0.677 | 16/29 | 0.151 | 0.564 | 5/9 |
| Full-Trajectory Guided Diffusion | 0.131 | 0.508 | 1/1 | 0.064 | 0.536 | 0/1 | 0.029 | 0.469 | 25/29 | 0.051 | 0.384 | 6/9 |
| siDiff w/o clustering | 0.016 | 0.371 | 1/1 | 0.059 | 0.516 | 0/1 | 0.028 | 0.478 | 21/29 | 0.026 | 0.513 | 6/9 |
| siDiff | 0.242 | 0.664 | 1/1 | 0.185 | 0.612 | 1/1 | 0.200 | 0.654 | 28/29 | 0.101 | 0.679 | 9/9 |

## Conclusion

In this paper, we presented an efficacy-guided discrete diffusion framework for de novo siRNA generation. Unlike conventional rank-based prediction models, our generative approach effectively balances sequence accuracy and structural diversity. By integrating target-aware biological constraints with an efficacy-guided sampling strategy, our method consistently outperforms state-of-the-art baselines across multiple benchmarks. Extensive ablation studies further validate the necessity and effectiveness of our guidance design. Notably, our model demonstrates superior generalization on real-world patent datasets, addressing a critical bottleneck in early-stage RNAi drug discovery.

## Supplementary Material Data curation

To enable robust validation across diverse data distributions, we constructed three novel datasets curated from patent repositories. However, a challenge with existing patent data is that *in vitro* screening is typically performed on chemically modified siRNAs. While modifications are essential for preserving structural stability and avoiding immune activation *in vivo*, they complicate the data. Consequently, the reported final inhibition rate is a joint function of both the naked siRNA sequence (which dictates target binding affinity) and the specific chemical modification pattern applied.

Let *x*_*i*_ denote the *i*-th naked siRNA sequence and *y*_*j*_ denote the *j*-th chemical modification pattern. The observed inhibition rate reported in the patent data can be formalized as:

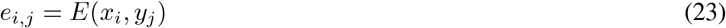

where *E* represents the underlying efficacy function.

To construct a dataset focused solely on the inherent binding efficacy of the naked sequences, we must decouple the sequence effect from the modification effect. Theoretically, while chemical modifications enhance systemic stability, they often introduce steric hindrance or thermodynamic changes that slightly attenuate the fundamental silencing efficacy compared to the unmodified sequence. Therefore, we assume that the true efficacy of the naked sequence, denoted as 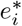, establishes an upper bound for any of its modified variants:

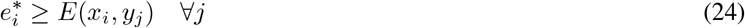

Because the true naked efficacy 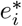 is rarely reported directly in these records, we approximate it using the available data. Let *y*_*i*_ = {*y*_*k*_} represent the empirical subset of modification patterns applied to sequence *x*_*i*_ within our dataset. We define our proxy label for the naked siRNA efficacy, 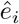, as the maximum observed efficacy across this sampled modification space:

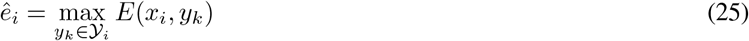

We focus our evaluation on three curated patent datasets targeting ANGPTL7, CTNNB1, and GSK3A, selected for their extensive experimental validation across diverse mRNA regions. By applying our maximum efficacy approximation to aggregate identical binding sites, the CTNNB1 dataset is distilled from 944 initial sequence-modification combinations (*x*_*i*_, *y*_*k*_) down to 330 unique naked sequences (*x*_*i*_). In contrast, for the ANGPTL7 and GSK3A datasets, all candidate siRNAs within a given target were evaluated using a uniform chemical modification pattern, denoted as (*x*_*i*_, *y*_0_). Because *y*_0_ is fixed for these targets, the modification space *y*_*i*_ consists of only a single pattern, allowing the reported inhibitions to map directly to our naked sequence proxy.

### Notation and Complete Pipeline Overview

This section follows the notation introduced in the main paper and provides a complete overview of the siDiff inference pipeline. We particularly clarify the distinction between the partial active pattern produced by guided diffusion and the complete siRNA candidates returned by the full pipeline.

### Notation

Let

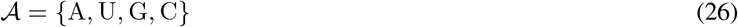

denote the nucleotide alphabet, and let

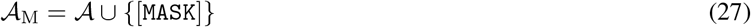

denote the augmented vocabulary used by the masked discrete diffusion model.

Following the problem formulation in the main paper, we use

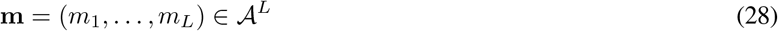

to denote an mRNA sequence or context supplied to the model.

A clean siRNA sequence is denoted by

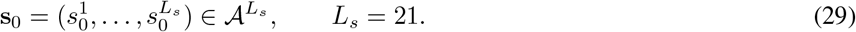

The partially masked state at diffusion timestep *t* is denoted by

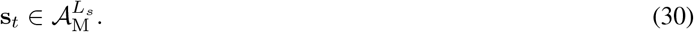

The forward diffusion process terminates at the fully masked state

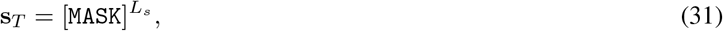

where *T* is the diffusion horizon.

In the training objective, the set of masked positions is

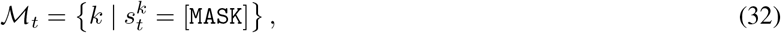

consistent with Eq. (6) of the main paper. During guided inference, we use

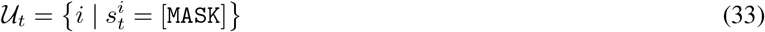

to denote the current set of unresolved positions. The two sets have the same token-level definition but are used in the training and sampling contexts, respectively.

The coarse-to-fine sampling process is controlled by two gates:

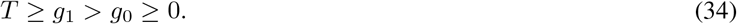

Active-pattern sampling is performed for

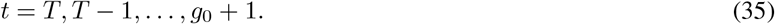

Position-level efficacy guidance is controlled by *w*_pos_(*t*), whereas base-level efficacy guidance is controlled by *β*_base_(*t*), which is disabled for *t > g*_1_ and enabled for *g*_0_ *< t* ≤ *g*_1_.

At timestep *t*, the conditional diffusion generator produces

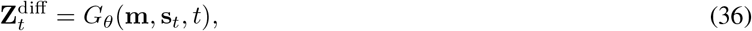

where

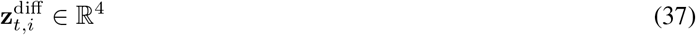

denotes the four nucleotide logits at position *i*. For every unresolved position *i* ∈ *U*_*t*_, the generation confidence is

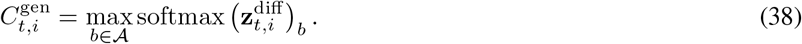

The partial-state efficacy predictor in Algorithm 1 of the main paper is denoted by *F*_*ϕ*_. For every unresolved position *i* and hypothetical nucleotide *b* ∈ A, it computes

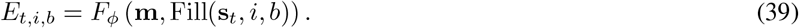

The position-level efficacy potential is

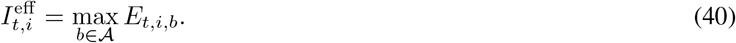

The four scores {*E*_*t,i,b*_} _*b* ∈ *A*_ are cached within each timestep and shared by position-level and base-level guidance. Table 6 summarizes the principal notation.

**Table 6:** Principal notation used in the supplementary material.

| Symbol | Definition |
| --- | --- |
| $\mathcal{A}$ | Nucleotide alphabet $\{A, U, G, C\}$ |
| $\mathcal{A}_M$ | Augmented alphabet containing [MASK] |
| $\mathbf{m}$ | mRNA sequence or context supplied to the model |
| $\mathbf{s}_0$ | Clean, complete 21-nt siRNA sequence |
| $\mathbf{s}_t$ | Partially masked siRNA state at timestep $t$ |
| $\mathcal{M}_t$ | Masked positions used in the diffusion training objective |
| $\mathcal{U}_t$ | Unresolved positions during guided sampling |
| $T$ | Diffusion horizon |
| $g_1$ | Timestep at which base-level guidance becomes active |
| $g_0$ | Terminal boundary of active-pattern sampling |
| $G_\theta$ | Conditional masked diffusion generator |
| $F_\phi$ | Partial-state efficacy predictor used during guided sampling |
| $\mathbf{z}_{t,i}^{\text{diff}}$ | Diffusion logits over the four nucleotides at position $i$ |
| $C_{t,i}^{\text{gen}}$ | Diffusion generation confidence at position $i$ |
| $E_{t,i,b}$ | Predicted efficacy after filling position $i$ with nucleotide $b$ |
| $I_{t,i}^{\text{eff}}$ | Position-level efficacy potential $\max_b E_{t,i,b}$ |
| $\tilde{C}_{t,i}$ | Modified position confidence after efficacy guidance |
| $w_{\text{pos}}(t)$ | Position-level efficacy-guidance weight |
| $\beta_{\text{base}}(t)$ | Base-level efficacy-guidance weight |
| $j_t$ | Position selected for unmasking at timestep $t$ |
| $\mathbf{s}^{\text{act}}$ | Partial active-pattern proposal produced by guided diffusion |
| $\mathcal{S}_{\text{cand}}$ | Set of complete siRNA candidates derived from $\mathbf{s}^{\text{act}}$ |
| $\mathcal{G}_k$ | The $k$-th cluster of candidate target sites |
| $q_a$ | Predicted efficacy score assigned to complete candidate $a$ |
| $\omega$ | Warm-up proportion of the position-guidance schedule |
| $\tau_{\text{pos}}$ | Temperature used by Gumbel-Top-1 position selection |
| $\Delta_{\text{clu}}$ | Maximum candidate-cluster span, defined as $L_s - g_0$ |

### Complete Pipeline Overview

The complete siDiff inference pipeline consists of three functional components: efficacy-guided active-pattern proposal, candidate derivation and clustering, and inhibition-aware candidate evaluation and sampling. Importantly, the partial sequence produced by guided diffusion is not directly returned as the final siRNA. Instead, it is used as an active pattern for proposing target regions. Every complete siRNA returned by the pipeline remains the reverse complement of a valid 21-nt mRNA target site.

### Efficacy-Guided Active-Pattern Proposal Starting from

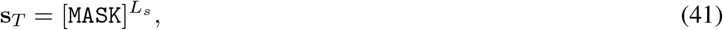

siDiff resolves one position at each reverse timestep. The diffusion generator first computes 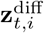 and 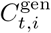 for all *i* ∈ *U*_*t*_. The efficacy predictor then evaluates the four hypothetical nucleotide assignments in Eq. (39) and computes 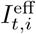.

Generation confidence and efficacy potential are independently normalized over the current unresolved-position set. Following Eq. (11) of the main paper, the modified position confidence is

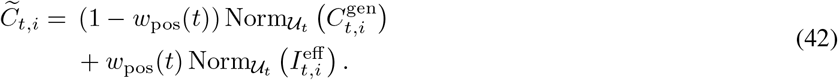

The position-guidance weight follows a quadratic progressive schedule. We first define the overall sampling progress as

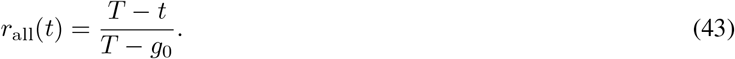

Let *ω* ∈ [0, 1) denote the warm-up proportion. The normalized post-warm-up progress is

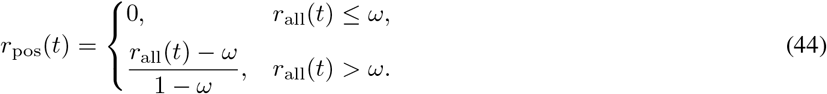

The normalized position-guidance schedule is

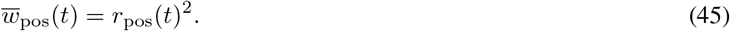

The implementation scales this normalized schedule by the configured position-guidance strength. Thus, position guidance is inactive during warm-up and increases quadratically afterward.

To retain stochastic exploration, the next position is selected using the Gumbel-Top-1 rule:

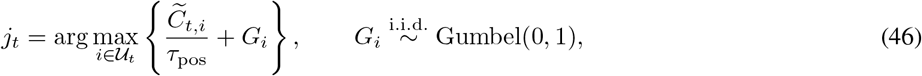

where *τ*_pos_ *>* 0 controls the exploration strength. A standard Gumbel random variable can be sampled as

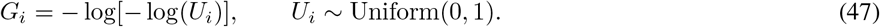

Its cumulative distribution function is

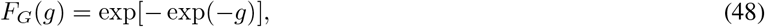

and its probability density function is

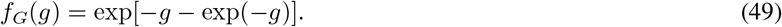

The Gumbel distribution is the Type-I limiting distribution for maxima of independent random variables under standard regularity conditions. This extreme-value property makes it suitable for stochastic Top-1 selection. By the Gumbel-Max identity,

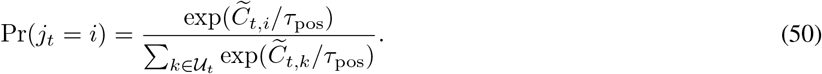

Thus, positions with larger modified confidence are more likely to be selected, while the Gumbel perturbation maintains stochastic exploration over different unmasking orders. For *t > g*_1_, the nucleotide at *j*_*t*_ is sampled using only the diffusion logits:

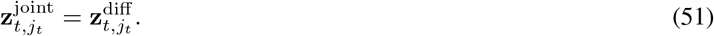

For *g*_0_ *< t* ≤ *g*_1_, the cached four-nucleotide efficacy scores are standardized:

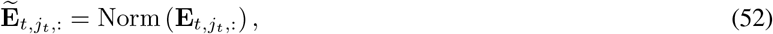

and incorporated into the nucleotide logits:

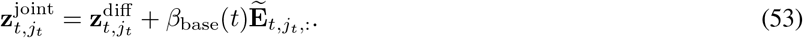

The normalized base-guidance schedule is

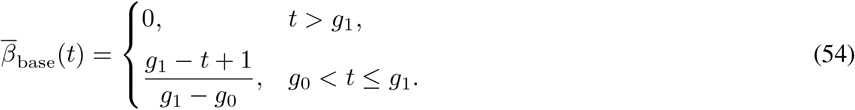

The implementation scales this normalized schedule by the configured base-guidance strength. Base-level guidance is therefore disabled for *t > g*_1_ and increases linearly from *t* = *g*_1_ to the final active-pattern sampling step *t* = *g*_0_ + 1. The nucleotide is then decoded from 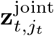. After the final sampling step *t* = *g*_0_ + 1, the remaining partial sequence is retained as the active pattern **s**^act^.

#### Candidate Derivation and Clustering

The active span of **s**^act^ is determined by its leftmost and rightmost instantiated nucleotides. Instantiated nucleotides are aligned to the target mRNA under reverse complementarity, while the remaining [MASK] positions are treated as unconstrained.

Matches satisfying the predefined mismatch tolerance are expanded into valid 21-nt mRNA target windows. Each target window is then converted into a complete siRNA through reverse complementation, yielding the complete candidate set *S*_cand_.

Candidates are sorted according to their target start positions and greedily grouped into spatial clusters. Following Eq. (17) of the main paper, a candidate with start position *x* is assigned to the current cluster C if

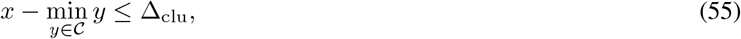

where

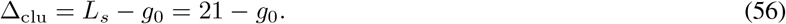

This choice connects the maximum cluster span to the number of positions resolved during active-pattern sampling. Otherwise, the candidate initializes a new cluster. Bounding the total cluster span preserves local alternatives while preventing spatially distant candidates from being merged through transitive chaining.

#### Inhibition-Aware Candidate Evaluation and Sampling

Following the terminology of the main paper, we refer to the inhibition predictor used in the *Inhibition-Aware Candidate Evaluation and Sampling* subsection as the full-sequence predictor. Unlike the partial-state efficacy predictor *F*_*ϕ*_, this predictor is applied after candidate derivation and clustering. It takes a clean, fully specified mRNA-context–siRNA pair as input and assigns a predicted efficacy score *q*_*a*_ to each complete candidate *a* ∈ *S*_cand_.

For every cluster *G*_*k*_, the mean predicted efficacy and high-efficacy ratio are

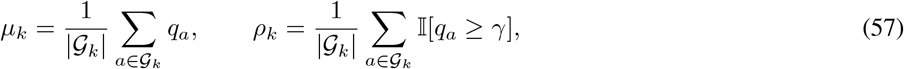

consistent with Eq. (18) of the main paper.

The three-grade staircase strategy assigns progressively smaller candidate quotas

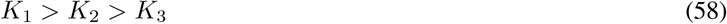

to clusters satisfying progressively relaxed thresholds on *µ*_*k*_ and *ρ*_*k*_. Clusters that fail the final grade are removed. Within each retained cluster, candidates are ranked by *q*_*a*_ and selected according to the assigned quota. This produces a final candidate set that balances predicted inhibition efficacy with coverage across distinct mRNA regions.

Algorithm 2 summarizes the complete inference pipeline.

##### Algorithm 2

Complete siDiff Inference Pipeline

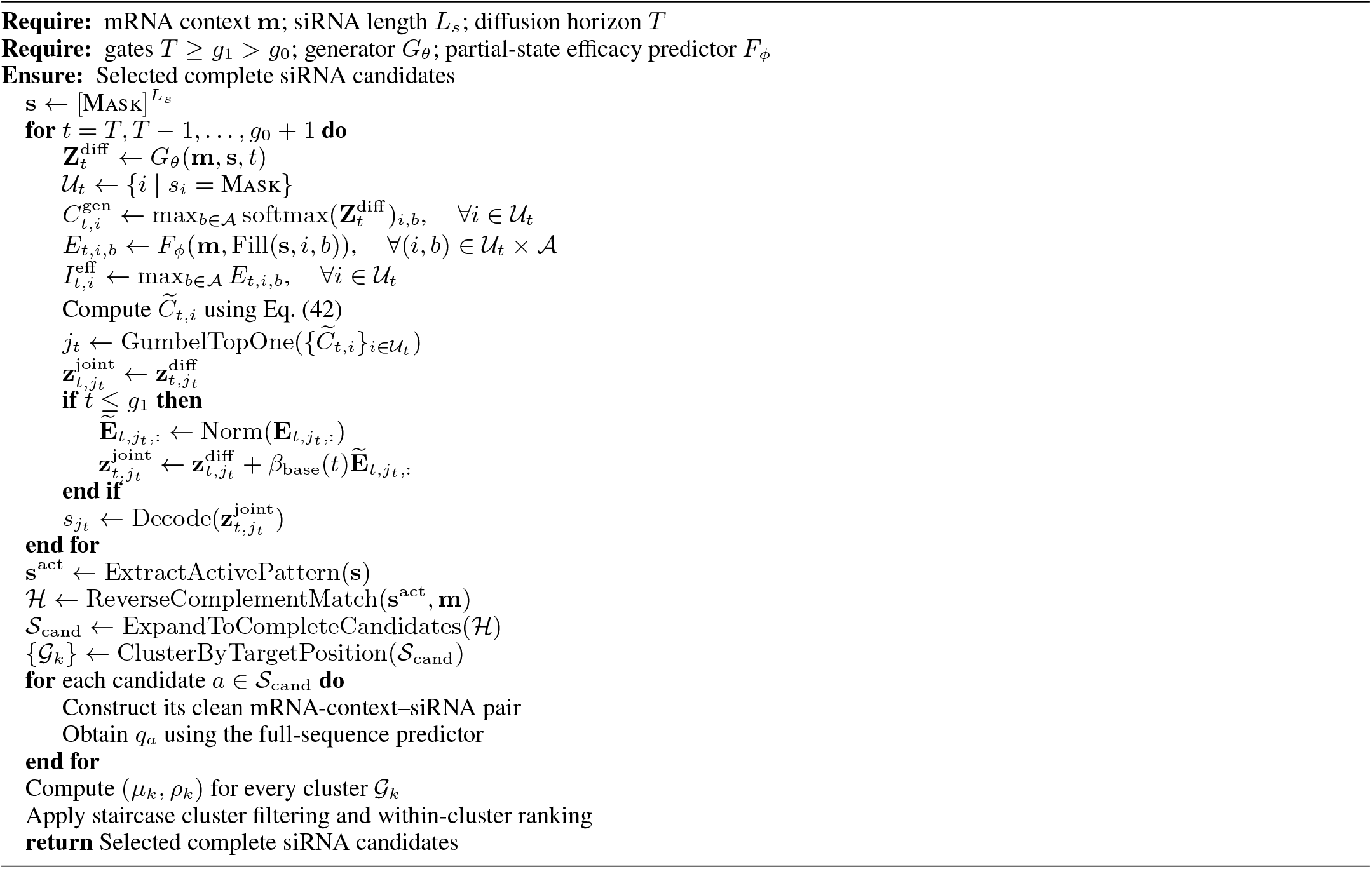

### Detailed Definition of Efficacy-Window Coverage

Efficacy-Window Coverage (EWC) measures whether a method recovers spatially diverse regions containing experimentally validated high-efficacy siRNA sites. For each evaluation mRNA, we partition its sequence into non-overlapping windows of width *w*. Specifically, for an mRNA of length *L*, the *j*-th window is

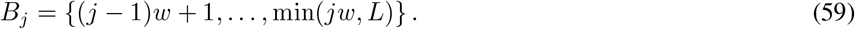

Let *O*_*j*_ denote the set of experimentally measured siRNA target sites whose start positions fall within *B*_*j*_, and let *y*_*i*_ ∈ [0, 1] be the experimentally measured efficacy of site *i* after per-mRNA min–max normalization. For each window, we compute

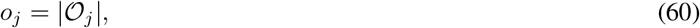

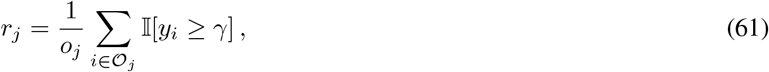

and

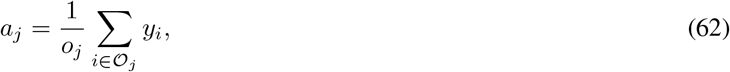

where *o*_*j*_, *r*_*j*_, and *a*_*j*_ respectively denote the number of measured sites, the proportion of high-efficacy sites, and the average measured efficacy within the window. A site is regarded as high efficacy when its normalized experimental efficacy satisfies *y*_*i*_ ≥ *γ*.

The ground-truth high-efficacy window set is defined as

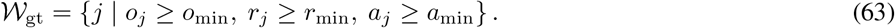

The fixed parameter configuration used for all four external evaluation datasets is

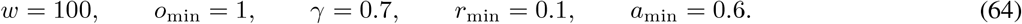

After deduplicating the target sites produced by a method, we construct its predicted high-efficacy window set *W*_pred_ analogously, replacing the experimental efficacy values with the efficacy scores produced by that method. Each method therefore ranks and filters its own candidates using its own output scores; scores from different predictors are neither pooled nor directly compared.

Finally, EWC is computed as

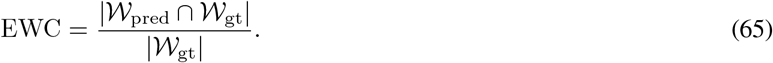

We report EWC in the count form |*W*_pred_ ∩ *W*_gt_| */* |*W*_gt_|.

### Efficacy Predictors Used at Different Stages

Our framework employs two efficacy predictors with distinct roles and input representations. The first predictor provides lightweight guidance over partially masked siRNA states during the reverse diffusion process, whereas the second evaluates complete candidates after candidate derivation and clustering. Both predictors are trained on the Huesken dataset, but they are trained separately and are not used interchangeably.

### Partial-State Efficacy Predictor for Guided Sampling

The partial-state efficacy predictor *F*_*ϕ*_ serves as the guidance model during reverse diffusion. Its input consists of a 21-nt mRNA target segment and a 21-nt siRNA state. To make the model applicable to intermediate diffusion states, partially masked siRNA inputs are constructed from the clean siRNAs during training, while the corresponding experimentally measured inhibition values are retained as supervision.

During generation, *F*_*ϕ*_ is repeatedly applied to the current partially decoded siRNA state to estimate the efficacy changes associated with candidate position and nucleotide assignments. We therefore use the 21-nt target segment without additional flanking context, which keeps the guidance model lightweight and makes repeated predictor evaluations during sampling computationally affordable.

### Full-Sequence Predictor for Candidate Evaluation

After candidate derivation and clustering, complete candidates are evaluated using the full-sequence predictor used in the *Inhibition-Aware Candidate Evaluation and Sampling* subsection of the main paper. This predictor takes a clean and complete mRNA-context–siRNA pair as input. Specifically, the mRNA context and siRNA have lengths of 31 and 21 nucleotides, respectively. The 31-nt mRNA input retains limited sequence context surrounding the target site while conforming to the fixed input-window length of 61 used by the predictor.

This predictor is also trained on the Huesken dataset. For each complete candidate *a*, it produces the efficacy score *q*_*a*_, which is subsequently used for candidate ranking and sampling within each cluster. In contrast to *F*_*ϕ*_, this predictor is applied only to the substantially reduced candidate set obtained after clustering and can therefore use the more contextualized input representation.

#### Distinction between the two predictors

The two predictors operate at different stages of the pipeline: *F*_*ϕ*_ guides local decisions over partially masked states, whereas the full-sequence predictor evaluates clean candidates using additional mRNA context. Their output scores are used only within their respective stages and are not directly compared across the two models.

### Masked Diffusion siRNA Model Training

#### Training data

The diffusion generator was trained exclusively on the Huesken-derived dataset. As described in the *Datasets* paragraph of the main paper, we retained siRNAs with normalized inhibition values of at least 0.7 and applied random-window context augmentation while preserving the complete 21-nt target site within each 61-nt mRNA context. This procedure produced 3,860 mRNA-context–siRNA training instances. We randomly split these instances into training and validation sets using an 80%*/*20% ratio, resulting in 3,088 training instances and 772 validation instances. The Takayuki and patent datasets (ANGPTL7, CTNNB1, and GSK3A) were not used during training or hyperparameter selection.

#### Generator architecture

The generator was implemented as a six-layer CrossAttentionDiT with a hidden dimension of 256 and eight attention heads. Each Transformer block consists of self-attention over the siRNA representation, cross-attention to the mRNA representation, and a feed-forward network. Layer normalization and timestep-dependent modulation are applied within each block. The diffusion model contains 7.6M trainable parameters.

#### Optimization

The generator was trained for 50 epochs using AdamW with a batch size of 64, a learning rate of 1 × 10^−4^, and a weight decay of 0.01. The learning rate was kept fixed throughout training. We used a cosine masking schedule to construct the corrupted diffusion states.

## Additional Experimental Results

### Reliability of the Partial-State Efficacy Predictor

We train *F*_*ϕ*_ using a curriculum that progressively expands the masking range from clean sequences to increasingly incomplete partial states. Table 7 compares its validation performance across curriculum stages and maximum mask levels.

**Table 7:** Validation performance of the partial-state efficacy predictor under different maximum masking levels and curriculum stages. Stage 1 is trained on clean sequences, Stage 2 introduces up to six masked nucleotides, and Stage 3 expands the training range to max masked number nucleotides.

| Max mask | Curriculum stage | PCC | SCC |
| --- | --- | --- | --- |
| 10 | Stage 1 | 0.628 | 0.624 |
|  | Stage 2 | 0.638 | 0.645 |
|  | Stage 3 | 0.653 | 0.660 |
| 12 | Stage 1 | 0.643 | 0.641 |
|  | Stage 2 | 0.663 | 0.669 |
|  | Stage 3 | **0.677** | **0.684** |
| 14 | Stage 1 | 0.675 | 0.686 |
|  | Stage 2 | 0.639 | 0.627 |
|  | Stage 3 | 0.601 | 0.624 |

The predictor maintains meaningful efficacy-ranking capability under progressively more challenging masking conditions. These results support the use of curriculum training to construct a mask-robust guidance model for intermediate diffusion states. Based on Table 7, we adopt the predictor trained through the full curriculum with a maximum mask level of 12 as the default guidance model *F*_*ϕ*_. This choice balances robustness to substantially incomplete siRNA states with the observation that further expanding the masking ceiling does not improve, and may even degrade, ranking performance. The curriculum comparison evaluates separately trained predictors under different masking ceilings, whereas the exact-mask evaluation fixes the selected max_m_ask = 12 checkpoint and varies only the number of masked positions. As shown in Table 8, *F*_*ϕ*_ maintains stable efficacy-ranking performance within its trained range (mask ≤ 12), while retaining meaningful rank correlation at adjacent mask counts beyond the curriculum ceiling. These results support selecting max_m_ask = 12 as a scientifically justified default for mask-robust guidance.

**Table 8:** Validation ranking performance of the default partial-state efficacy predictor (max_mask = 12, Stage 3) under exact mask counts.

| Mask count | Status | PCC | SCC |
| --- | --- | --- | --- |
| 11 | In range | $0.674 \pm 0.003$ | $0.687 \pm 0.004$ |
| 12 | In range | $0.681 \pm 0.002$ | $0.690 \pm 0.002$ |
| 13 | Adjacent | $0.617 \pm 0.008$ | $0.671 \pm 0.004$ |
| 14 | Adjacent | $0.643 \pm 0.007$ | $0.659 \pm 0.005$ |

### Sensitivity to the Guidance Gates

The gates *g*_0_ and *g*_1_ control two complementary aspects of guided sampling. We evaluate different combinations of the two gates in Table 9.

**Table 9:**
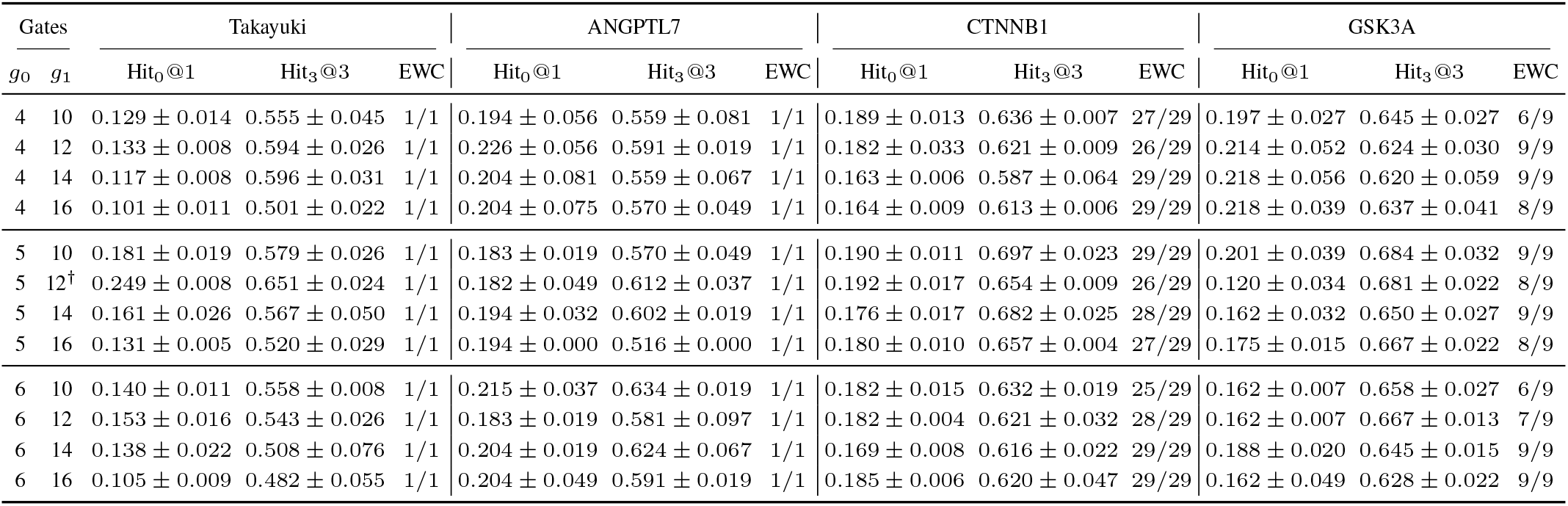
Sensitivity to the guidance gates *g*_0_ and *g*_1_ on four datasets.

| Gates |  | Takayuki |  |  | ANGPTL7 |  |  | CTNNB1 |  |  | GSK3A |  |  |
| --- | --- | --- | --- | --- | --- | --- | --- | --- | --- | --- | --- | --- | --- |
| $g_0$ | $g_1$ | Hit<sub>0</sub>@1 | Hit<sub>3</sub>@3 | EWC | Hit<sub>0</sub>@1 | Hit<sub>3</sub>@3 | EWC | Hit<sub>0</sub>@1 | Hit<sub>3</sub>@3 | EWC | Hit<sub>0</sub>@1 | Hit<sub>3</sub>@3 | EWC |
| 4 | 10 | 0.129 ± 0.014 | 0.555 ± 0.045 | 1/1 | 0.194 ± 0.056 | 0.559 ± 0.081 | 1/1 | 0.189 ± 0.013 | 0.636 ± 0.007 | 27/29 | 0.197 ± 0.027 | 0.645 ± 0.027 | 6/9 |
| 4 | 12 | 0.133 ± 0.008 | 0.594 ± 0.026 | 1/1 | 0.226 ± 0.056 | 0.591 ± 0.019 | 1/1 | 0.182 ± 0.033 | 0.621 ± 0.009 | 26/29 | 0.214 ± 0.052 | 0.624 ± 0.030 | 9/9 |
| 4 | 14 | 0.117 ± 0.008 | 0.596 ± 0.031 | 1/1 | 0.204 ± 0.081 | 0.559 ± 0.067 | 1/1 | 0.163 ± 0.006 | 0.587 ± 0.064 | 29/29 | 0.218 ± 0.056 | 0.620 ± 0.059 | 9/9 |
| 4 | 16 | 0.101 ± 0.011 | 0.501 ± 0.022 | 1/1 | 0.204 ± 0.075 | 0.570 ± 0.049 | 1/1 | 0.164 ± 0.009 | 0.613 ± 0.006 | 29/29 | 0.218 ± 0.039 | 0.637 ± 0.041 | 8/9 |
| 5 | 10 | 0.181 ± 0.019 | 0.579 ± 0.026 | 1/1 | 0.183 ± 0.019 | 0.570 ± 0.049 | 1/1 | 0.190 ± 0.011 | 0.697 ± 0.023 | 29/29 | 0.201 ± 0.039 | 0.684 ± 0.032 | 9/9 |
| 5 | 12<sup>†</sup> | 0.249 ± 0.008 | 0.651 ± 0.024 | 1/1 | 0.182 ± 0.049 | 0.612 ± 0.037 | 1/1 | 0.192 ± 0.017 | 0.654 ± 0.009 | 26/29 | 0.120 ± 0.034 | 0.681 ± 0.022 | 8/9 |
| 5 | 14 | 0.161 ± 0.026 | 0.567 ± 0.050 | 1/1 | 0.194 ± 0.032 | 0.602 ± 0.019 | 1/1 | 0.176 ± 0.017 | 0.682 ± 0.025 | 28/29 | 0.162 ± 0.032 | 0.650 ± 0.027 | 9/9 |
| 5 | 16 | 0.131 ± 0.005 | 0.520 ± 0.029 | 1/1 | 0.194 ± 0.000 | 0.516 ± 0.000 | 1/1 | 0.180 ± 0.010 | 0.657 ± 0.004 | 27/29 | 0.175 ± 0.015 | 0.667 ± 0.022 | 8/9 |
| 6 | 10 | 0.140 ± 0.011 | 0.558 ± 0.008 | 1/1 | 0.215 ± 0.037 | 0.634 ± 0.019 | 1/1 | 0.182 ± 0.015 | 0.632 ± 0.019 | 25/29 | 0.162 ± 0.007 | 0.658 ± 0.027 | 6/9 |
| 6 | 12 | 0.153 ± 0.016 | 0.543 ± 0.026 | 1/1 | 0.183 ± 0.019 | 0.581 ± 0.097 | 1/1 | 0.182 ± 0.004 | 0.621 ± 0.032 | 28/29 | 0.162 ± 0.007 | 0.667 ± 0.013 | 7/9 |
| 6 | 14 | 0.138 ± 0.022 | 0.508 ± 0.076 | 1/1 | 0.204 ± 0.019 | 0.624 ± 0.067 | 1/1 | 0.169 ± 0.008 | 0.616 ± 0.022 | 29/29 | 0.188 ± 0.020 | 0.645 ± 0.015 | 9/9 |
| 6 | 16 | 0.105 ± 0.009 | 0.482 ± 0.055 | 1/1 | 0.204 ± 0.049 | 0.591 ± 0.019 | 1/1 | 0.185 ± 0.006 | 0.620 ± 0.047 | 29/29 | 0.162 ± 0.049 | 0.628 ± 0.022 | 9/9 |

The results show that the method remains effective across a range of gate configurations. Default configuration is 5 and 12 for *g*_0_ and *g*_1_, respectively.

### Comparison of Guidance Modes

To examine whether the sampling behavior depends on a particular guidance schedule, we compare several position- and nucleotide-level guidance variants while keeping the generator, efficacy predictor, gate configuration, and candidate-processing pipeline unchanged. The default configuration uses quadratic position guidance with a maximum weight of 0.5 and linear nucleotide-level guidance with a maximum weight of 1.0. For position guidance, we additionally test a constant schedule with weight 0.5, a linear schedule with maximum weight 0.5, and a stronger quadratic schedule whose maximum weight is increased to 1.0. For nucleotide-level guidance, we compare the default linear schedule against a constant schedule with weight 1.0, a quadratic schedule with maximum weight 1.0 and exponent 2, and a stronger linear schedule with maximum weight 2.0. Each variant modifies only one guidance component, while the other remains at its default setting. The results are summarized in Table 10.

**Table 10:** Comparison of guidance-mode variants on four external datasets. All variants use *g*_0_ = 5 and *g*_1_ = 12.

| Guidance mode | Takayuki |  |  | ANGPTL7 |  |  | CTNNB1 |  |  | GSK3A |  |  |
| --- | --- | --- | --- | --- | --- | --- | --- | --- | --- | --- | --- | --- |
|  | Hit<sub>0</sub>@1 | Hit<sub>3</sub>@3 | EWC | Hit<sub>0</sub>@1 | Hit<sub>3</sub>@3 | EWC | Hit<sub>0</sub>@1 | Hit<sub>3</sub>@3 | EWC | Hit<sub>0</sub>@1 | Hit<sub>3</sub>@3 | EWC |
| Position-Constant | 0.135 | 0.586 | 1/1 | 0.226 | 0.677 | 1/1 | 0.178 | 0.645 | 28/29 | 0.205 | 0.679 | 8/9 |
| Position-Linear | 0.226 | 0.620 | 1/1 | 0.194 | 0.613 | 1/1 | 0.190 | 0.657 | 27/29 | 0.167 | 0.590 | 8/9 |
| Position-Larger Max | 0.181 | 0.595 | 1/1 | 0.159 | 0.616 | 1/1 | 0.190 | 0.653 | 29/29 | 0.244 | 0.679 | 9/9 |
| Base-Constant | 0.089 | 0.537 | 1/1 | 0.194 | 0.581 | 1/1 | 0.161 | 0.653 | 26/29 | 0.192 | 0.667 | 7/9 |
| Base-Quadratic | 0.137 | 0.552 | 1/1 | 0.194 | 0.581 | 1/1 | 0.161 | 0.641 | 29/29 | 0.180 | 0.679 | 9/9 |
| Base-Larger Max | 0.216 | 0.540 | 1/1 | 0.194 | 0.563 | 1/1 | 0.194 | 0.636 | 29/29 | 0.154 | 0.628 | 9/9 |
| Position-Quadratic, Base-Linear | 0.242 | 0.664 | 1/1 | 0.185 | 0.612 | 1/1 | 0.200 | 0.654 | 28/29 | 0.101 | 0.679 | 9/9 |

Overall, the default combination of quadratic position guidance and linear nucleotide-level guidance provides the most stable trade-off between preserving the learned generative prior, introducing efficacy preference, and maintaining candidate coverage. Constant schedules apply guidance uniformly throughout the corresponding sampling phase, whereas linear and quadratic schedules progressively strengthen the guidance signal. Increasing the maximum weight further emphasizes efficacy-driven selection, but may reduce the balance between guided optimization and generative exploration.

### Comparison with Enumeration-Based Methods

Enumeration-based approaches exhaustively construct valid siRNAs and then rely on a predictor to rank the resulting candidates. Their ability to identify experimentally potent siRNAs is therefore directly limited by the predictor’s generalization to previously unseen targets. To examine this limitation, we compare the scores produced by the full-sequence predictor used in the *Inhibition-Aware Candidate Evaluation and Sampling* subsection of the main paper, OligoFormer, and iScore against experimentally measured inhibition values. The results are reported in Table 11.

**Table 11:** Agreement between predictor scores and experimentally measured inhibition on four datasets. The full-sequence predictor refers to the predictor used in the *Inhibition-Aware Candidate Evaluation and Sampling* subsection of the main paper. PCC and SCC measure point-wise linear and rank correlations, respectively. Lower RMSE is better.

| Predictor | Takayuki |  |  | ANGPTL7 |  |  | CTNNB1 |  |  | GSK3A |  |  |
| --- | --- | --- | --- | --- | --- | --- | --- | --- | --- | --- | --- | --- |
|  | PCC↑ | SCC↑ | RMSE↓ | PCC↑ | SCC↑ | RMSE↓ | PCC↑ | SCC↑ | RMSE↓ | PCC↑ | SCC↑ | RMSE↓ |
| Full-sequence predictor (ours) | 0.079 | 0.087 | 0.270 | 0.164 | 0.141 | 0.290 | 0.167 | 0.187 | 0.328 | 0.054 | 0.129 | 0.345 |
| OligoFormer | 0.458 | 0.482 | 0.195 | 0.383 | 0.407 | 0.243 | 0.055 | 0.082 | 0.302 | 0.268 | 0.247 | 0.297 |
| iScore | 0.452 | 0.467 | 0.193 | 0.292 | 0.292 | 0.250 | 0.029 | 0.042 | 0.315 | 0.229 | 0.185 | 0.308 |

The predictors exhibit substantial variation across external datasets, and strong performance on one target does not consistently transfer to other targets. These results demonstrate that exhaustive enumeration alone cannot resolve out-of-distribution predictor errors, motivating siDiff’s combination of generative proposal, efficacy-guided sampling, and structured candidate selection.

### Regional Analysis of Efficacy-Window Coverage

To further analyze Efficacy-Window Coverage, we visualize the regional agreement between experimentally defined high-efficacy windows and those recovered by siDiff by partitioning each full-length mRNA into contiguous 100-nt windows. A window is considered high efficacy if it contains at least one measured site, at least 10% of its measured sites have efficacy greater than 0.7, and its mean efficacy is at least 0.6. Figures 3–6 show the window-wise mean efficacy of the ground-truth and siDiff-recovered high-efficacy windows for Takayuki, ANGPTL7, CTNNB1, and GSK3A, with non-high-efficacy windows displayed as zero. Across these transcripts, siDiff achieves high regional recall while accurately localizing the experimentally identified high-efficacy regions. Moreover, the number of predicted high-efficacy windows remains close to that of the ground-truth set, indicating that the improved recall is not obtained through indiscriminate overprediction.

**Figure 3.**
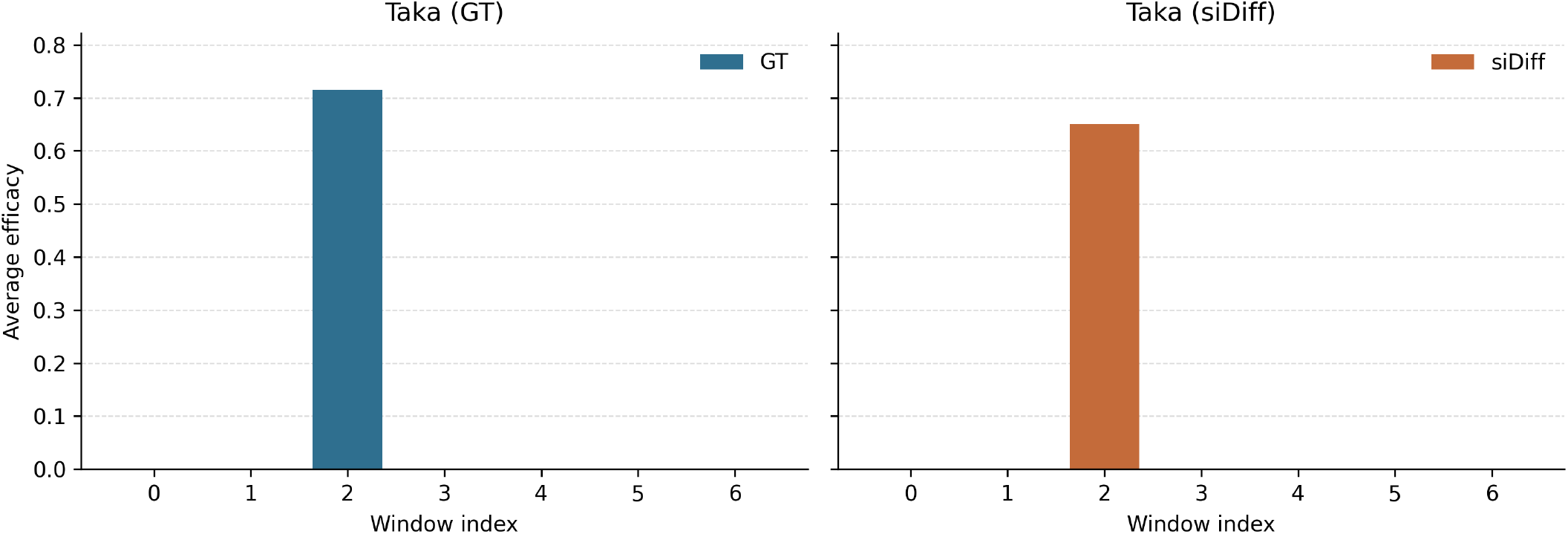
Ground-truth and siDiff high-efficacy windows on Takayuki.

**Figure 4.**
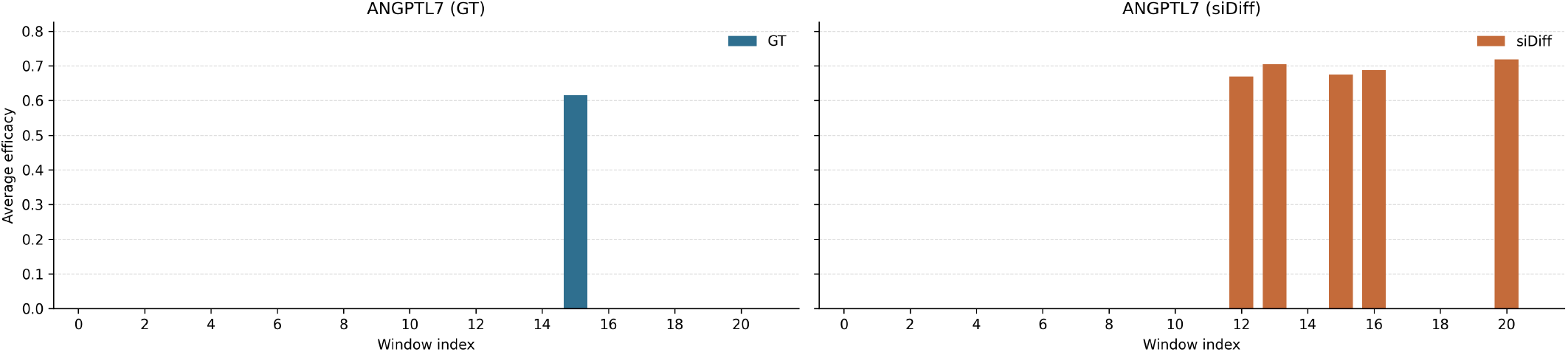
Ground-truth and siDiff high-efficacy windows on ANGPTL7.

**Figure 5.**
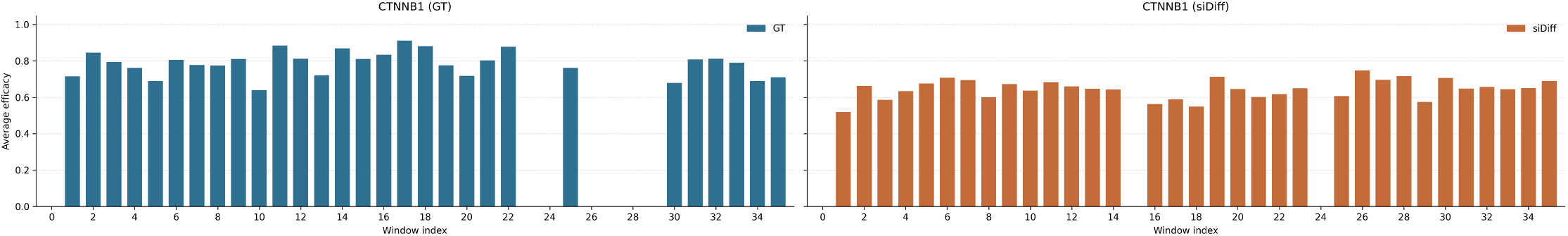
Ground-truth and siDiff high-efficacy windows on CTNNB1.

**Figure 6.**
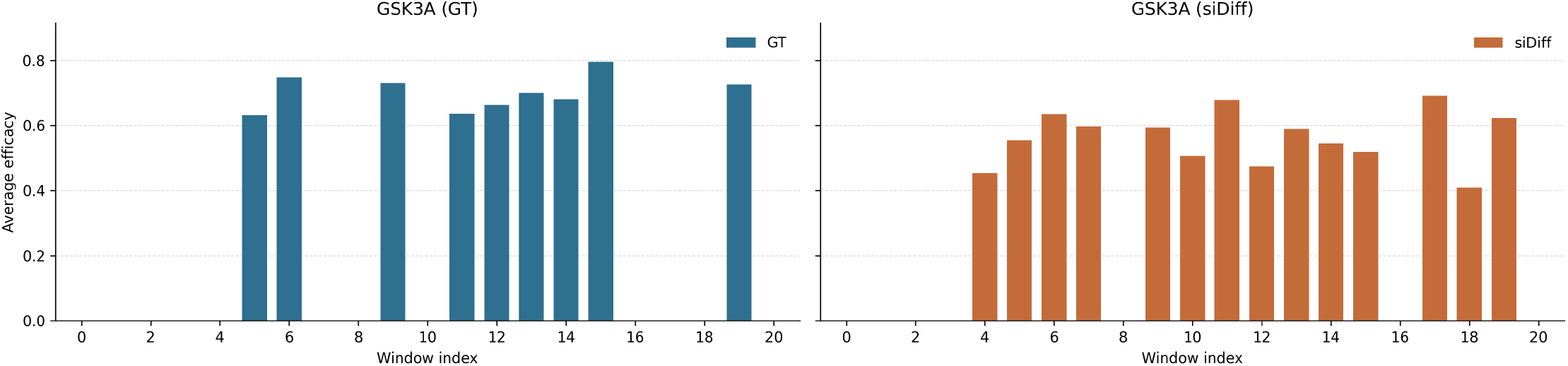
Ground-truth and siDiff high-efficacy windows on GSK3A.

